# Increased mitochondrial activity upon CatSper channel activation is required for sperm capacitation

**DOI:** 10.1101/2021.05.04.442630

**Authors:** Juan J. Ferreira, Adriana Cassina, Pilar Irigoyen, Mariana Ford, Santiago Pietroroia, Rafael Radi, Celia M. Santi, Rossana Sapiro

## Abstract

To fertilize an oocyte, sperm must become hyperactive. However, whether they obtain ATP for hyperactivated motility via glycolysis or mitochondrial respiration is unclear. Here, high-resolution respirometry, flow cytometry, and confocal microscopy experiments revealed that mitochondrial respiration and membrane potential increased during mouse sperm capacitation. Treatment with inhibitors of mitochondrial respiration prevented sperm from hyperactivating and fertilizing an oocyte. Mitochondrial respiration was impaired in sperm from mice lacking the calcium channel CatSper. We developed a method to image mitochondrial calcium in sperm and found that CatSper activation led to increased mitochondrial calcium concentration. Finally, treating sperm with an inhibitor of mitochondrial calcium import impaired mitochondrial function and sperm hyperactivation. Together, our results uncover a new role of sperm mitochondria and reveal a new pathway connecting calcium influx through CatSper to mitochondrial activity and the sperm hyperactivation required to fertilize an oocyte.

**Summary:** The source of ATP for sperm hyperactivation is unclear. Ferreira et al. show that mitochondrial activity increases during, and is required for, hyperactivation and fertilization ability. Increased mitochondrial activity depends on calcium influx through the channel CatSper.

## Introduction

To fertilize an oocyte, sperm must undergo several biochemical and functional changes known as capacitation. During capacitation, sperm switch from progressive to hyperactivated motility and undergo a regulated release of acrosomal content in a process called the acrosome reaction (AR) (1–3). These processes, which normally occur in the female reproductive tract, allow sperm to free themselves from the oviduct wall, penetrate the zona pellucida, and fuse with the oocyte. An important event controlling capacitation is an increase in intracellular [Ca^2+^] (4, 5) mediated by the cation sperm-specific channel (CatSper) (6–8). *CatSper* Knockout (KO) mice are infertile because their sperm fail to hyperactivate and fertilize oocytes. However, the downstream effects of CatSper-mediated Ca^2+^ increase in sperm have not been fully elucidated (9, 10).

In other cell types, regulated Ca^2+^ entry into mitochondria increases the efficiency of oxidative respiration (11, 12) via activation of many Ca^2+^-dependent mitochondrial enzymes resulting in the increase of ATP production (13). Whether Ca^2+^ plays similar roles in sperm mitochondria is unknown (14). Complicating this picture is the unclear role of mitochondria in sperm capacitation. Although some data suggest that sperm rely on glycolysis instead of mitochondrial respiration to produce ATP for motility and fertilization (15), other studies showed that mitochondrial respiration increases during capacitation in both human (16–18) and mouse (19) sperm. However, the significance of this increase in mitochondrial activity during capacitation and its role in sperm hyperactivation, acrosome reaction (AR), and fertilization remains to be established.

Given that sperm from CatSper KO mice have reduced ATP production (20), we hypothesized that Ca^2+^ influx through CatSper channels during capacitation enhances mitochondrial activity, thereby contributing to sperm hyperactivation and fertilization. To test our hypothesis, we studied mitochondrial activity using high-resolution respirometry (HRR), measurements of mitochondrial membrane potential (MMP) and evaluation of the ATP/ADP exchange during capacitation, as well as measurements of mitochondrial Ca^2+^, both in wild-type and CatSper KO sperm. In addition, we analyzed the effects of mitochondrial function inhibitors on hallmark parameters of capacitation: AR, hyperactivation, tyrosine phosphorylation and the ability of the sperm to fertilize the egg.

## Results

### Mitochondrial respiration increases in capacitated mouse sperm

High-resolution respirometry (HRR) (21) was performed in non-capacitated (NC) and capacitated (CAP) sperm. **Figure 1A** shows a representative trace of oxygen consumption rate measured in capacitated sperm. The mean respiratory control ratio (RCR) (5.01 ± 1.0 vs. 3.2 ± 0.3) and coupling efficiency (0.52 ± 0.04 vs. 0.43 ± 0.03) were both significantly higher in CAP than in NC sperm (**Figure 1B** and **Table 1**). We found no significant difference in spare respiratory capacity. We next measured MMP in NC or CAP sperm with the lipophilic cationic dye tetramethyl rhodamine methyl ester perchlorate (TMRM) (22–24). By flow cytometry we detected two sperm populations (M1 and M2, **Figure 1C**) with different MMP in both NC and CAP sperm. Similar populations have been reported previously in human sperm (23). We found that the normalized fluorescence of the M2 population was significantly higher in CAP sperm than in NC sperm (13.22 ± 6.6 vs. 8.83 ± 7.80, n=8, **Figure 1D**). In addition, the M2 peak was completely abolished upon addition of the uncoupling agent carbonyl cyanide 4-(trifluoromethoxy) phenylhydrazone (FCCP) in both NC and CAP sperm. These data suggest that MMP in the M2 population increases during capacitation. We also used confocal imaging to measure TMRM fluorescence in the midpiece of individual NC and CAP sperm (see **Figure 1E, F** for a representative example), we found that TMRM fluorescence was significantly higher in CAP than in NC sperm (3.27 ± 0.50 vs. 2.05 ± 0.36, **Figure 1G**). Taken together, these data reveal that mitochondrial activity is higher CAP than in NC sperm.

**Figure 1.**
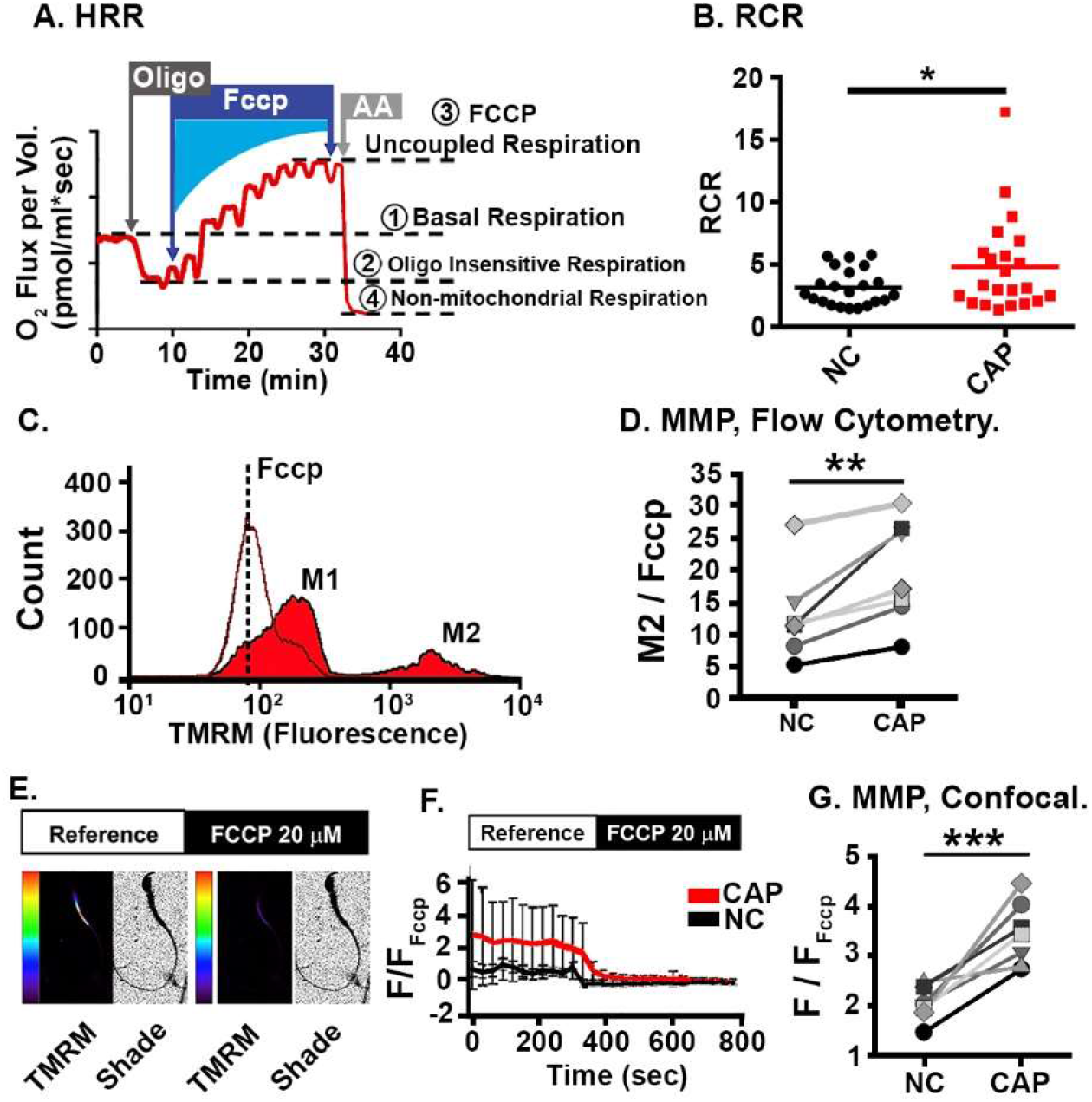
Capacitated sperm exhibit increased mitochondrial function. **A.** Representative trace of oxygen consumption rate (red line) from CAP mouse sperm. Sperm cells were exposed sequentially to oligomycin (oligo), Carbonyl cyanide 4-(trifluoromethoxy) phenylhydrazone (FCCP), and Antimycin A (AA). Oxygen consumption (QO2) levels used to calculate mitochondrial indexes are indicated by dotted lines and numbered from 1 to 4. **B**. Respiratory Control Ratio (RCR) values measured from wild-type CAP and NC sperm. Lines represent mean RCR values of NC (black dots) and CAP sperm (red dots) (n=18). **C.** Representative histograms of TMRM fluorescence from CAP sperm, recorded with flow cytometry. **D.** Graph shows normalized TMRM fluorescence values (MMP) of M2 cell population under NC and CAP conditions (n=7.) **E.** Representative confocal images of TMRM fluorescence from CAP control and FCCP-treated conditions. **F.** Paired representative normalized traces of TMRM fluorescence from NC and CAP sperm. Fluorescence was measured only in the sperm midpiece, and only cells responding to FCCP were included in the measurements. **G.** Graphs show normalized TMRM sperm midpiece fluorescence for NC and CAP sperm samples (n=8). For **B** and **D**. paired *t-test* was used to determine statistical significance and an independent *t-test* was used in G. * p < 0.05 and *** p <0.001.

**Table 1:**
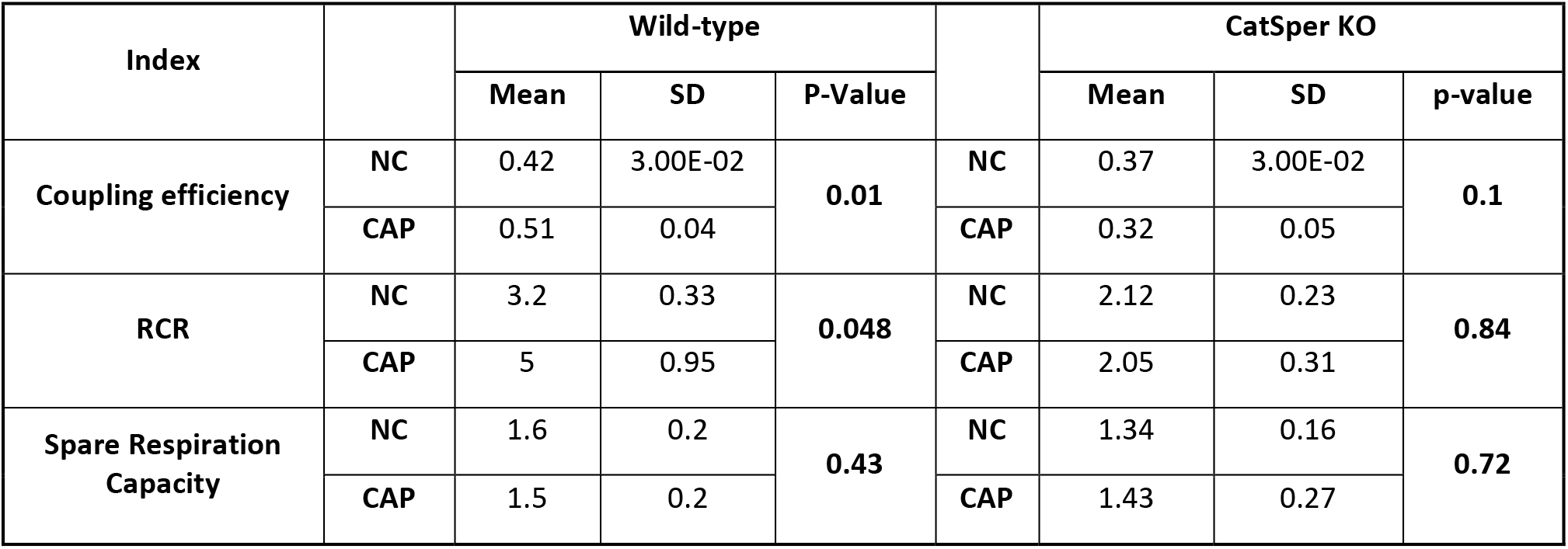
Oximetry values in NC and CAP sperm from wild-type and CatSper KO mice. Coupling efficiency= Oligomycin-sensitive respiration/basal respiration; RCR = Maximal respiration in the presence of FCCP/ATP turnover; spare respiratory capacity = Maximal respiration in the presence of FCCP/Basal Respiration.

| Index |  | Wild-type |  |  |  | CatSper KO |  |  |
| --- | --- | --- | --- | --- | --- | --- | --- | --- |
|  |  | Mean | SD | P-Value |  | Mean | SD | p-value |
| Coupling efficiency | NC | 0.42 | 3.00E-02 | 0.01 | NC | 0.37 | 3.00E-02 | 0.1 |
|  | CAP | 0.51 | 0.04 |  | CAP | 0.32 | 0.05 |  |
| RCR | NC | 3.2 | 0.33 | 0.048 | NC | 2.12 | 0.23 | 0.84 |
|  | CAP | 5 | 0.95 |  | CAP | 2.05 | 0.31 |  |
| Spare Respiration Capacity | NC | 1.6 | 0.2 | 0.43 | NC | 1.34 | 0.16 | 0.72 |
|  | CAP | 1.5 | 0.2 |  | CAP | 1.43 | 0.27 |  |

### Mitochondrial respiration during capacitation is important for hyperactivated motility and oocyte fertilization

To address whether mitochondrial activity was required for the functional hallmarks of capacitation, we capacitated sperm for one hour in control conditions or in the presence of FCCP or AA. A significantly higher percentage of sperm were hyperactivated in control conditions than in the presence of FCCP or AA (20.43 ± 9.89, n=15, vs. 10.51 ± 8.44, n=15, vs. 7.58 ± 7.76, n=11, **Figure 2A**). In contrast, FCCP and AA had no effect on the percentage of sperm that underwent spontaneous or induced AR (**Figure 2B**). Additionally, AA had no effect on protein tyrosine phosphorylation (**Supplementary Figure 1**). Sperm treated with FCCP or AA were significantly less able to fertilize oocytes than untreated sperm (**Figure 2C and 2D**).

**Figure 2.**
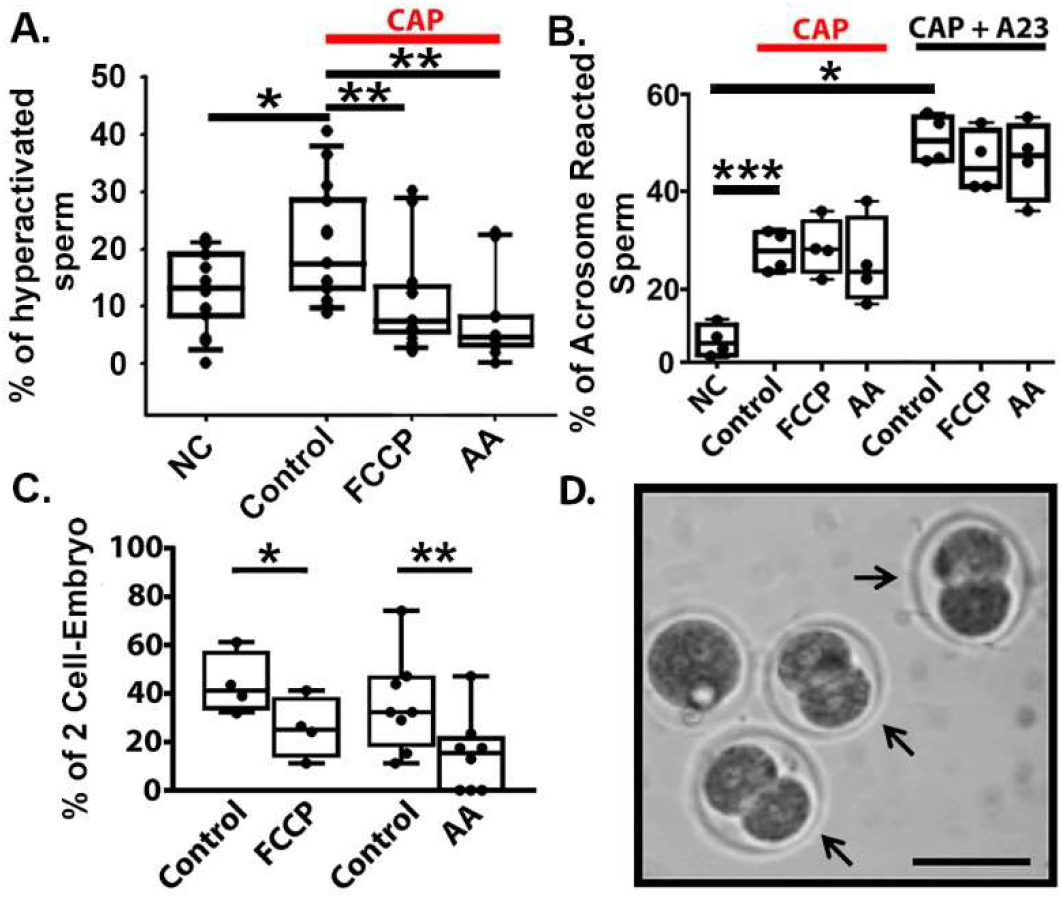
Mitochondrial inhibitors impair mouse sperm capacitation and *in vitro* fertilization. **A.** Hyperactivation measurements obtained by CASA from CAP sperm in control conditions and in the presence of FCCP and AA. **B.** Spontaneous AR and A23187-induced AR from NC, CAP control, and CAP with FCCP or AA. **C.** Percentage of oocytes that reached the 2-cells stage embryo after 24 hours of sperm addition. Graph shows mean ± SD. **D.** Example of the 2-cell stage (arrows) from C. Bar represents a 50 μm scale. To determine statistical significance between the groups, independent *t-test* were used on A and B, and Paired *t-test* was used in C. * p < 0.05, ** p < 0.01, and *** p < 0.001 respectively.

Together, these data indicate that while mitochondrial activity during capacitation is not required for the AR or tyrosine phosphorylation, it significantly contributes to sperm hyperactivation and, notably, to their ability to fertilize oocytes.

### Mitochondrial respiration is impaired in sperm from CatSper knockout mice

By analyzing HRR in sperm from CatSper KO mice we found that respiration values in sperm from CatSper KO mice did not differ between NC and CAP conditions (**Figure 3 A, B** and **Table 1**). The MMP showed no statistical difference between CAP and NC in CatSper KO mice (2.90 ± 0.92 vs. 3.10 ± 0.48, *P*=0.746, **Figure 3C, D**). **Figure 3E** shows the variation in MMP after CAP for wild-type and CatSper KO mice; MMP increase in wild-type sperm during capacitation is statistically larger than in the KO sperm. Our results indicate that CatSper is involved in increasing sperm mitochondrial activity during capacitation.

**Figure 3.**
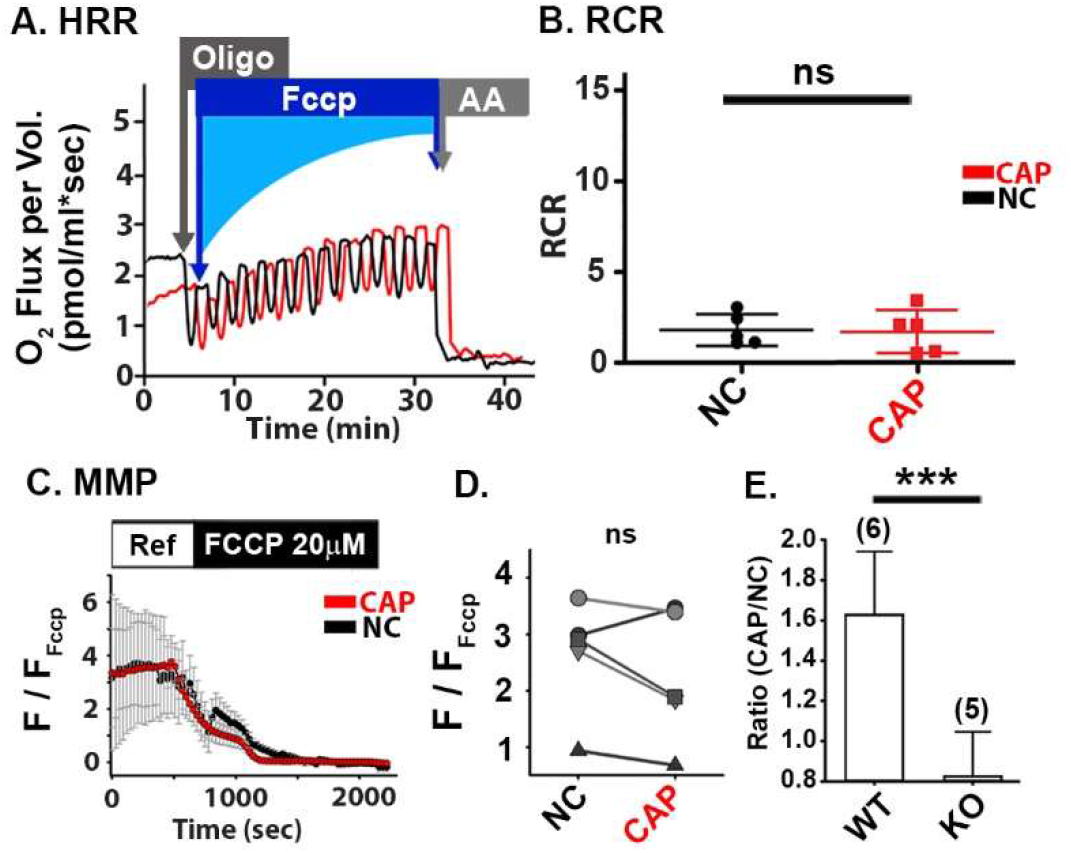
CatSper KO sperm lacks capacitation-induced mitochondrial function increase. **A.** Representative recordings of a HRR from CAP, and NC in CatSper KO sperm. Oxygen consumption was measured in control conditions and after the addition of mitochondrial inhibitors, Oligomycin, FCCP and AA. **B.** Respiratory control ratio (RCR) measurements from CAP and NC CatSper KO mouse sperm. A paired *t-test* was used to determine statistical significance (n=5). **C.** Paired representative traces of normalized TMRM fluorescence, under CAP and NC conditions. **D.** Graph shows normalized TMRM fluorescence under NC and CAP conditions (n=5). **E.** TMRM fluorescence ratio after CAP (normalized to NC) in wild-type and CatSper KO mice. To determine statistical significance, paired *t-test* was used. ns p > 0.050 and *** p < 0.001. Error bars represent SD.

### Mitochondrial Ca^2+^ concentration increases during capacitation in sperm from wild-type but not CatSper KO mice

We next wanted to test whether the CatSper-dependent increase in cytoplasmic [Ca^2+^] during capacitation leads to an increase in mitochondrial [Ca^2+^]. To do so, we developed a new method to measure mitochondrial [Ca^2+^] in mouse sperm by using the fluorescent Ca^2+^-indicator, Fluo-5N. Whereas Fluo-4 measures [Ca^2+^] in the 100 nM – 300 nM range **(Supplementary Figure 2)** and is suitable for measuring cytoplasmic [Ca^2+^], Fluo-5N has a lower Ca^2+^-binding affinity and is suitable for measuring [Ca^2+^] in the 1 μM to 1 mM range, appropriate for measuring [Ca^2+^] in mitochondria (25). We first confirmed that Fluo-5N co-localized in the sperm midpiece with mitochondrial markers. In sperm from Acr-eGFP+Su9-Red2 transgenic mice, which express green fluorescent protein (GFP) in the acrosome and red fluorescent protein in mitochondria (RFP), Fluo-5N co-localized with RFP in the sperm midpiece **(Supplementary Figure 3A)**. Likewise, Fluo-5N co-localized with MitoTracker^®^ red in the sperm midpiece **(Supplementary Figure 3B)**. In contrast, Fluo-4 distributed throughout the entire sperm **(Supplementary Figure 3C)**. Although Fluo-5N was also detected in the acrosome **(Supplementary Figure 3A)**, we excluded this in our assays by only measuring fluorescence in the sperm midpiece. To further confirm that our assay measured mitochondrial [Ca^2+^], we loaded sperm with either Fluo-5N or Fluo-4 and continuously measured fluorescence in the midpiece before and after adding FCCP **(Supplementary Figure 4A)**. As expected, Fluo-5N fluorescence decreased in the midpiece upon addition of FCCP **(Supplementary Figure 4B)**, whereas Fluo-4 fluorescence in the midpiece increased **(Supplementary Fig 4C)**, suggesting that FCCP caused Ca^2+^ to be released from the mitochondria into the cytosol. Thus, we concluded that our method accurately measured [Ca^2+^] of the mitochondria in the sperm midpiece.

We then applied this method to determine if mitochondrial Ca^2+^ content was different between NC and CAP wild-type sperm. **Figure 4A** shows representative images of CAP wild-type sperm loaded with Fluo-5N. At 2 mM extracellular [Ca^2+^], we observed that mitochondrial Ca^2+^ was higher in CAP than in NC wild-type sperm **(Figure 4B and D)**. In contrast, when we performed the experiment at 0 mM extracellular [Ca^2+^], we observed no difference in mitochondrial Ca^2+^ content between CAP and NC wild-type sperm **(Figure 4B and D)**. This is an expected result, as capacitation does not occur in the absence of extracellular calcium.

**Figure 4.**
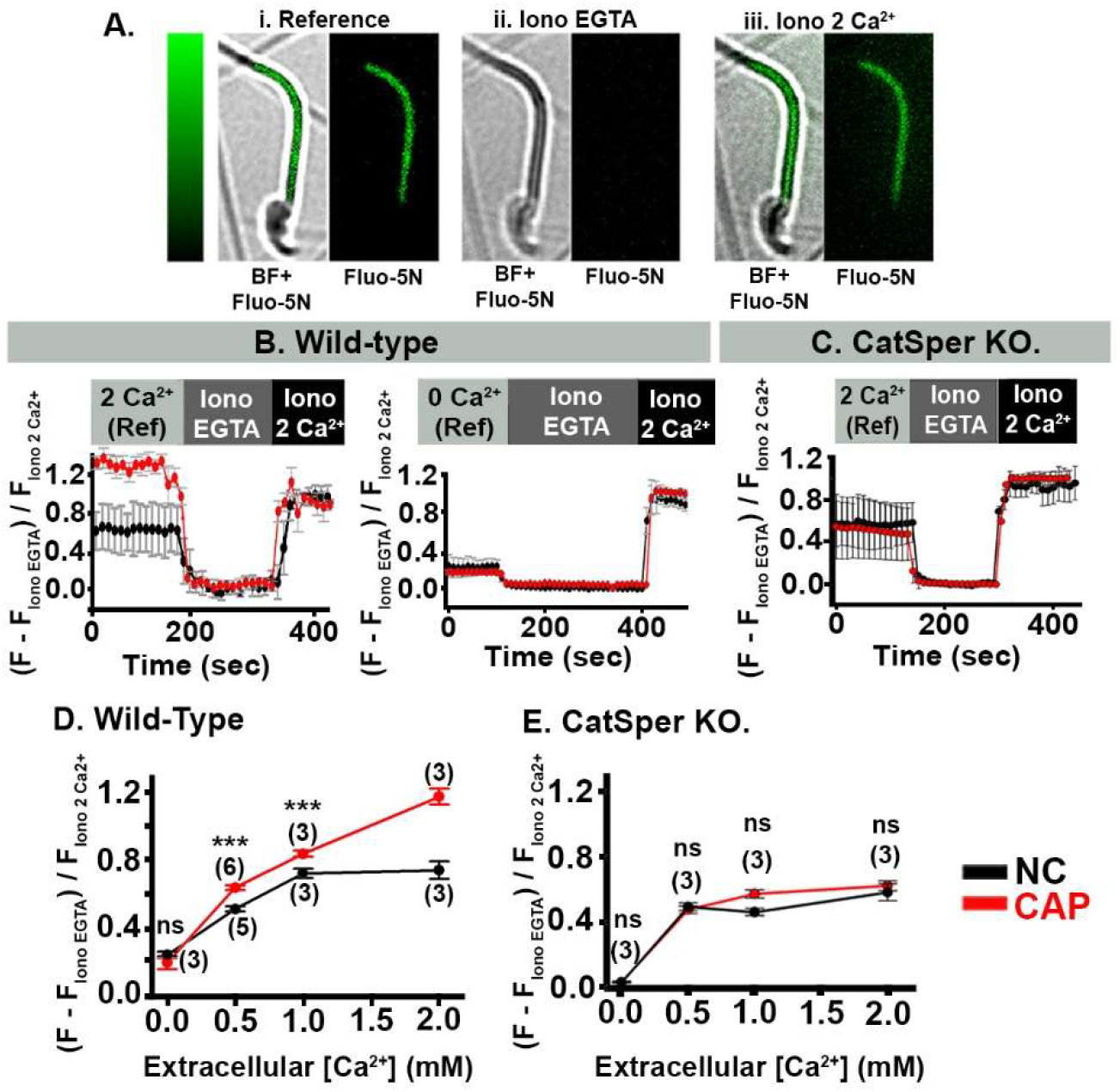
Sperm mitochondrial [Ca^2+^] increases during capacitation (CAP) in wild-type, but not in CatSper KO sperm. **A.** Representative images of CAP sperm loaded with Fluo-5N; and perfused with i) 2 mM Ca^2+^ (Reference), ii) 0 Ca^2+^ + 2 mM EGTA and Ionomycin (Iono EGTA), iii) Ionomycin + 2 mM Ca^2+^ (Iono 2 Ca^2+^). Images in i, ii, iii are on the left: bright field (BF) merged with Fluo-5N fluorescence (FITC). Right, Fluo-5N fluorescence. **B and C**, Representative traces of Fluo-5N fluorescence in the sperm midpiece from NC (black) and CAP (red) sperm incubated in 2 mM Ca^2+^ **(B, left)**, and in 0 mM Ca^2+^ **(B, right)**. **C.** Representative traces of Fluo-5N fluorescence from NC and CAP CatSper KO sperm. **D.** Normalized values of Fluo-5N fluorescence at reference (ref), in NC and CAP sperm at different extracellular [Ca^2+^]. **E.** Fluo-5N fluorescence at reference in CatSper KO samples, obtained under NC and CAP conditions, at different extracellular [Ca^2+^]. Statistical significance between CAP and NC samples at each extracellular [Ca^2+^] were evaluated with an independent *t-test.* Fluorescence values are shown in Table 2. ns p > 0.050, and *** p < 0.001.

We found that mitochondrial Ca^2+^ content increased with higher extracellular [Ca^2+^] (0, 0.5, 1 and 2 mM) in both NC and CAP wild-type sperm, however, the increase was significantly larger in CAP sperm **(Figure 4D, Table 2)**. Whereas mitochondrial Fluo-5N fluorescence reached a plateau at 1 mM extracellular Ca^2+^ in NC sperm, the Fluo-5N fluorescence further increased at 2 mM extracellular Ca^2+^ in CAP sperm, consistently with the extracellular [Ca^2+^] reported to achieve capacitation (26).

**Table 2.** Fluo-5N fluorescent values corresponding to mitochondrial [Ca^2+^] before and after CAP at different extracellular [Ca^2+^]. Fluo-5N fluorescence values were normalized using Ionomycin + 0 mM Ca^2+^ + EGTA (Iono EGTA) values and Ionomycin + 2 mM Ca^2+^ (Iono 2 Ca^2+^) values. To determine statistical significance between NC and CAP samples at each extracellular [Ca^2+^], independent *t-test* were used.

| Extracellular [Ca<sup>2+</sup>] mM |  | Wild-type |  |  | P-Value |  | CatSper KO |  |  | p-value |
| --- | --- | --- | --- | --- | --- | --- | --- | --- | --- | --- |
|  |  | Mean | SD | n |  |  | Mean | SD | n |  |
| 0 | NC | 0.2398 | 0.1556 | 3 | 0.197 | NC | 0.0201 | 0.0167 | 3 | 0.396 |
|  | CAP | 0.1935 | 0.1136 | 3 |  | CAP | 0.0234 | 0.0248 | 3 |  |
| 0.5 | NC | 0.5122 | 0.2798 | 5 | 0.035 | NC | 0.4893 | 0.1984 | 3 | 0.676 |
|  | CAP | 0.6394 | 0.2413 | 6 |  | CAP | 0.4717 | 0.1995 | 3 |  |
| 1 | NC | 0.7242 | 0.2798 | 3 | 0.01 | NC | 0.4562 | 0.1649 | 3 | 0.63 |
|  | CAP | 0.8402 | 0.208 | 3 |  | CAP | 0.5688 | 0.257 | 3 |  |
| 2 | NC | 0.7427 | 0.4494 | 3 | 0.022 | NC | 0.579 | 0.3106 | 3 | 0.155 |
|  | CAP | 1.1791 | 0.4635 | 3 |  | CAP | 0.6194 | 0.3805 | 3 |  |

We conclude that mitochondrial [Ca^2+^] depends on extracellular [Ca^2+^]. Importantly, mitochondrial [Ca^2+^] are significantly higher in CAP than in NC wild-type sperm. To determine whether the increase in mitochondrial [Ca^2+^] in CAP sperm was dependent on CatSper activity, we measured changes in mitochondrial [Ca^2+^] during capacitation in sperm from CatSper KO mice. Our results showed that in CatSper KO sperm, at 2 mM extracellular [Ca^2+^], there is no difference in Fluo-5N fluorescence at the midpiece between NC and CAP conditions **(Figure 4C)**. When we plotted values of fluorescence in the midpiece of sperm from CatSper KO mice, we found that mitochondrial [Ca^2+^] only increased when we changed extracellular [Ca^2+^] from 0 to 0.5 mM, and then it remained constant. This increase was similar in NC and CAP sperm **(Figure 4E)**. The fact that Ca^2+^ increased at all in response to increased extracellular [Ca^2+^] in sperm from CatSper KO mice suggests that Ca^2+^ can enter sperm through another pathway. Consistent with this idea, by using ratiometric measurements (Fura 2 AM), we found that sperm from CatSper KO mice contained a cytosolic [Ca^2+^] of 90-100 nM (data not shown). Nonetheless, mitochondrial [Ca^2+^] was significantly lower in CAP sperm from CatSper KO mice than in CAP sperm from wild-type mice **(Figure 4D, E and Table 2)**.

Another way to validate that CatSper channels were involved in the increased mitochondrial [Ca^2+^], was to treat sperm from wild-type mice with the CatSper channel blocker Mibefradil during capacitation. Mibefradil (20 μM) significantly reduced the increase of mitochondrial [Ca^2+^] associated with capacitation **(Supplementary Figure 5)**. Taken together these results confirmed that mitochondrial [Ca^2+^] increase involved CatSper activation.

**Figure 5.**
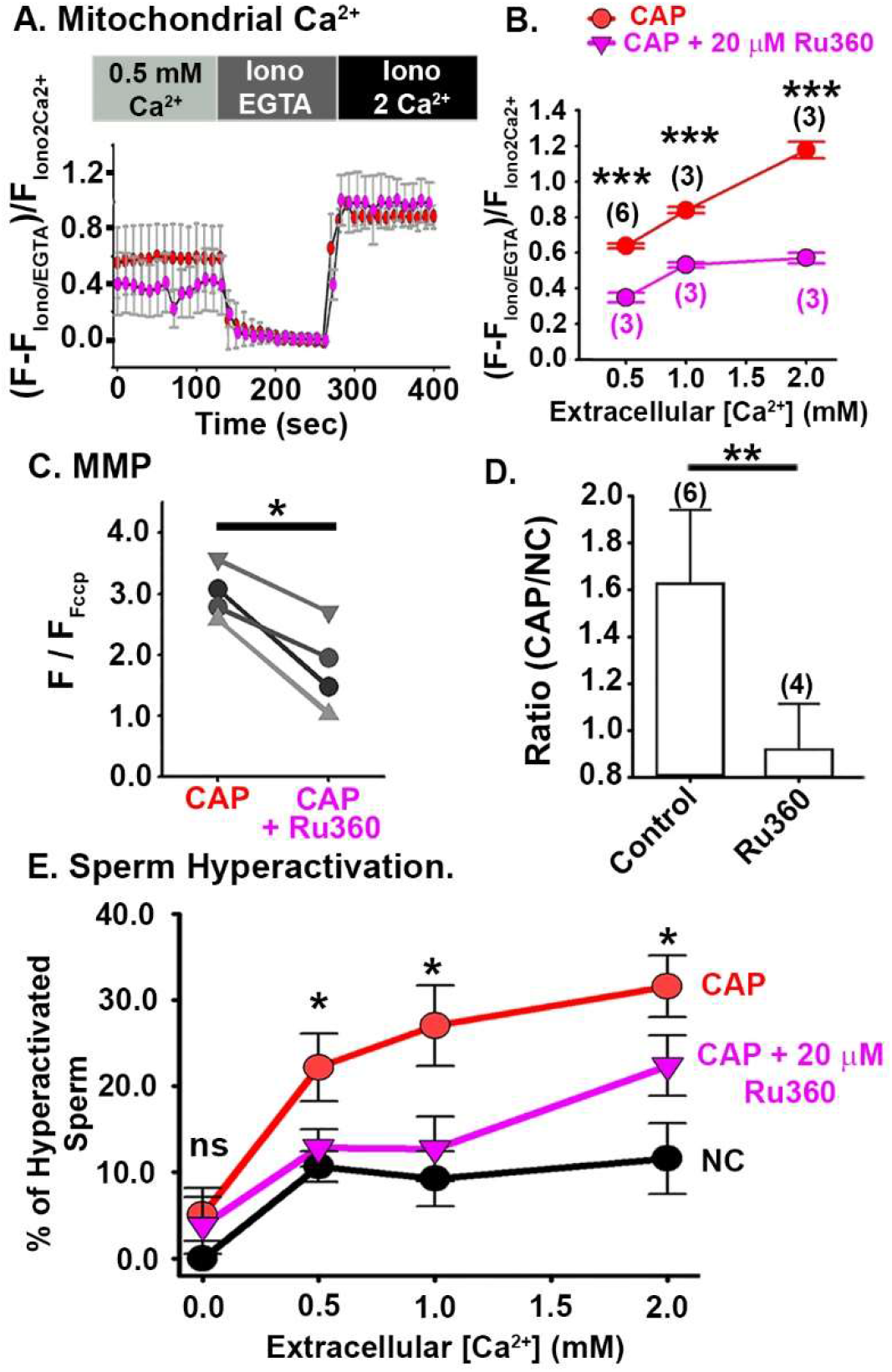
Effect of the Mitochondrial Calcium Uniporter inhibitor, Ru360 on mitochondrial [Ca^2+^], mitochondrial function, and sperm hyperactivation. **A.** Representative Fluo-5N fluorescence traces from CAP sperm and CAP with Ru360 at 0.5 mM extracellular Ca^2+^. **B.** Normalized Fluo-5N fluorescence in NC and CAP conditions, at different extracellular [Ca^2+^]. Graph shows mean values +/-SD of cells responding to Iono EGTA and Iono 2Ca^2+^. An independent *t-test* was used to determined statistical difference. **C.** Normalized TMRM fluorescence under CAP and CAP+Ru360 conditions (n=4). To determine statistical significance, paired *t-test* were used. **D.** TMRM fluorescence ratio in wild-type sperm, under CAP (control) and CAP+Ru360. To determine statistical significance an independent *t-test* was used. **E.** Percentage of hyperactivated sperm measured by CASA from sperm samples incubated in NC, CAP or CAP + 20 μM Ru360 media, at different extracellular [Ca^2+^]. Values are also presented in Table 4. An independent *t-test* were used to evaluate statistical significance between NC, CAP and CAP + Ru360 groups at different [Ca^2+^] (n=5). Error Bars in figure represent SD. ns P > 0.050, * p < 0.050, ** p<0.010, and *** p<0.001.

### Increase in mitochondrial function associated with capacitation depends on the mitochondrial Ca^2+^ uniporter (MCU)

We suspected that the increase of mitochondrial [Ca^2+^] in CAP sperm occurred via Ca^2+^ influx through the MCU, which resides in the internal mitochondrial membrane (27). To test this idea, we treated sperm with the specific MCU inhibitor Ru360 during capacitation (28) and then measured capacitation-associated changes in mitochondrial [Ca^2+^], MMP, and sperm hyperactivation.

First, mitochondrial [Ca^2+^] was significantly reduced after capacitation in the presence of Ru360 when compared with those without Ru360 **(Figure 5A-B and Table 3)**. Second, we treated CAP sperm with vehicle or Ru360, loaded them with TMRM in the presence of 2 mM extracellular Ca^2+^, and used our confocal microscopy assay to measure MMP. The normalized TMRM fluorescence was lower in sperm treated with Ru360 than in those without Ru360 (1.789 ± 0.717 vs. 3.00 ± 0.43, P = 0.027, **Figure 5C**). Moreover, whereas the MMP was 60% higher in CAP than NC sperm treated with vehicle, the MMP was 5% lower in CAP than NC sperm treated with Ru360 **(Figure 5D)**. Thus, we concluded that Ru360 abolished the MMP increase observed in wild-type sperm after capacitation.

**Table 3.** Fluo-5N fluorescence values corresponding to mitochondrial Ca^2+^ from wild-type sperm CAP in the presence of Ru360 at different extracellular [Ca^2+^]. Fluo-5N values were normalized using Ionomycin + 0 mM Ca^2+^ + EGTA (Iono EGTA) values and Ionomycin + 2 mM Ca^2+^ (Iono 2 Ca^2+^) values.

| Extracellular<br>[Ca<sup>2+</sup>] mM |  | Wild-type |  |  |
| --- | --- | --- | --- | --- |
|  |  | Mean | SD | n |
| 0.5 | CAP+Ru360 | 0.3483 | 0.027 | 3 |
| 1 | CAP+Ru360 | 0.5324 | 0.015 | 3 |
| 2 | CAP+Ru360 | 0.5725 | 0.0298 | 3 |

Finally, we assessed the effects of Ru360 on sperm hyperactivation by performing computer-assisted sperm analysis (CASA). In NC sperm, only 10% of sperm were hyperactive even at the highest extracellular [Ca^2+^] tested (2 mM). In CAP samples treated with vehicle, approximately 30% of sperm were hyperactive at 2 mM extracellular [Ca^2+^]. However, Ru360 treatment significantly diminished capacitation-induced hyperactivation **(Figure 5E and Table 4)**. To exclude the possibility that Ru360 impaired hyperactivation by directly inhibiting CatSper channels, we used Fluo-4 AM fluorescence to measure Ca^2+^ influx through CatSper channels in the absence and presence of Ru360. We found that membrane depolarization with KCl still triggers an increase of cytoplasmic [Ca^2+^] even in the presence of Ru360, confirming that this compound did not inhibit Ca^2+^ influx through CatSper channels **(Supplementary Figure 6A)**. Together, these results suggest that the increased in mitochondrial [Ca^2+^] contributes to sperm hyperactivation, and that this increase depends on extracellular [Ca^2+^].

**Table 4.** Percentage of hyperactivation from Figure 5E. Hyperactivated motility was measured with CASA in NC, CAP (Control), and CAP+20μM Ru360 samples at different [Ca^2+^]. Independent *t-test* was used to evaluate statistical significance between NC, CAP and CAP + Ru360 groups at the same [Ca^2+^].

| Extracellular<br>[Ca<sup>2+</sup>] mM |  | Wild-type |  |  | p-value |  |
| --- | --- | --- | --- | --- | --- | --- |
|  |  | Mean | SD | n |  |  |
| 0 | NC | 0 | 2.50E-03 | 4 | 0.197 |  |
|  | CAP | 5.0865 | 6.1481 | 4 |  | 0.91 |
|  | CAP + Ru360 | 3.8123 | 7.3331 | 5 |  |  |
| 0.5 | NC | 10.6899 | 4.3254 | 6 | 0.035 |  |
|  | CAP | 22.1885 | 9.6493 | 6 |  | 0.04 |
|  | CAP + Ru360 | 12.85 | 5.283 | 6 |  |  |
| 1 | NC | 9.2572 | 7.7815 | 6 | 0.01 |  |
|  | CAP | 25.8553 | 10.9269 | 7 |  | 0.03 |
|  | CAP + Ru360 | 17.4244 | 7.8503 | 7 |  |  |
| 2 | NC | 11.5959 | 9.2466 | 5 | 0.022 |  |
|  | CAP | 28.3665 | 10.678 | 6 |  | 0.03 |
|  | CAP + Ru360 | 17.7953 | 10.2253 | 6 |  |  |

### Hyperactivation relies on ADP/ATP translocase activity

In **Figure 1B**, we showed that the RCR was higher in CAP sperm than in NC sperm, suggesting that mitochondria in CAP sperm produce more ATP, which could be required for sperm hyperactivation. To test this idea, we inhibited ATP/ADP exchange between the mitochondria and the cytosol by capacitating sperm in the presence of Atractyloside (ATR), which inhibits the adenine nucleotide translocase residing in the inner mitochondrial membrane. Measurements with CASA, showed that hyperactivation was significantly lower in sperm treated with ATR than in those treated with vehicle (26.57 ± 5.95, n= 10, vs. 34.02 ± 8.77, n=9, P<0.01, **Figure 6A**). To exclude the possibility that ATR impaired hyperactivation by directly inhibiting CatSper, we used Fluo-4 AM fluorescence to measure Ca^2+^ influx through CatSper in the absence and presence of ATR. This experiment confirmed that ATR did not inhibit Ca^2+^ influx through CatSper **(Supplementary Figure 6B)**. When we compared the percentages of inhibited hyperactivation in the presence of Ru360 or ATR, the values were similar (35.77 ± 26.85, n= 6, vs. 26.42 ± 10.41, n=6; **Figure 6B**). This result suggests that Ca^2+^ entry into mitochondria during capacitation is important for the ATP production involved in sperm hyperactivation.

**Figure 6.**
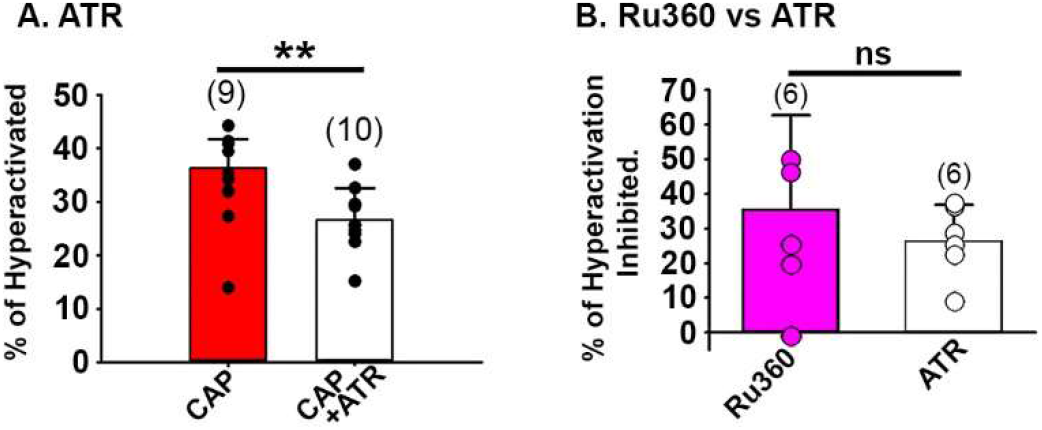
Mouse sperm hyperactivation is reduced by ADP/ATP translocase inhibitor Atractyloside (ATR). **A.** Percentage of hyperactivated sperm measured by CASA from CAP Control and CAP + 5 μM ATR conditions. Measurements were done in media with 2 mM extracellular Ca^2+^. Independent *t-test* was used to evaluate statistical significance between CAP and CAP+ATR. Error Bars represent SD. **B**. Graph shows percentage of hyperactivation inhibited by the presence of Ru360 or ATR during capacitation. An independent *t-test* was used to evaluate statistical significance. Error Bars represent SD. ns p > 0.050, and ** p < 0.010.

## Discussion

Together, our data support the following model **(Figure 7):** CatSper channel activation during sperm capacitation facilitates Ca^2+^ influx, thereby increasing cytoplasmic [Ca^2+^]. This increase in intracellular [Ca^2+^] starts in the principal piece, propagates through the midpiece, and reaches the head in a few seconds (20). The MCU then transports Ca^2+^ into the mitochondria, leading to increased mitochondrial efficiency, which promotes sperm hyperactivation and their ability to fertilize an oocyte.

**Figure 7.**
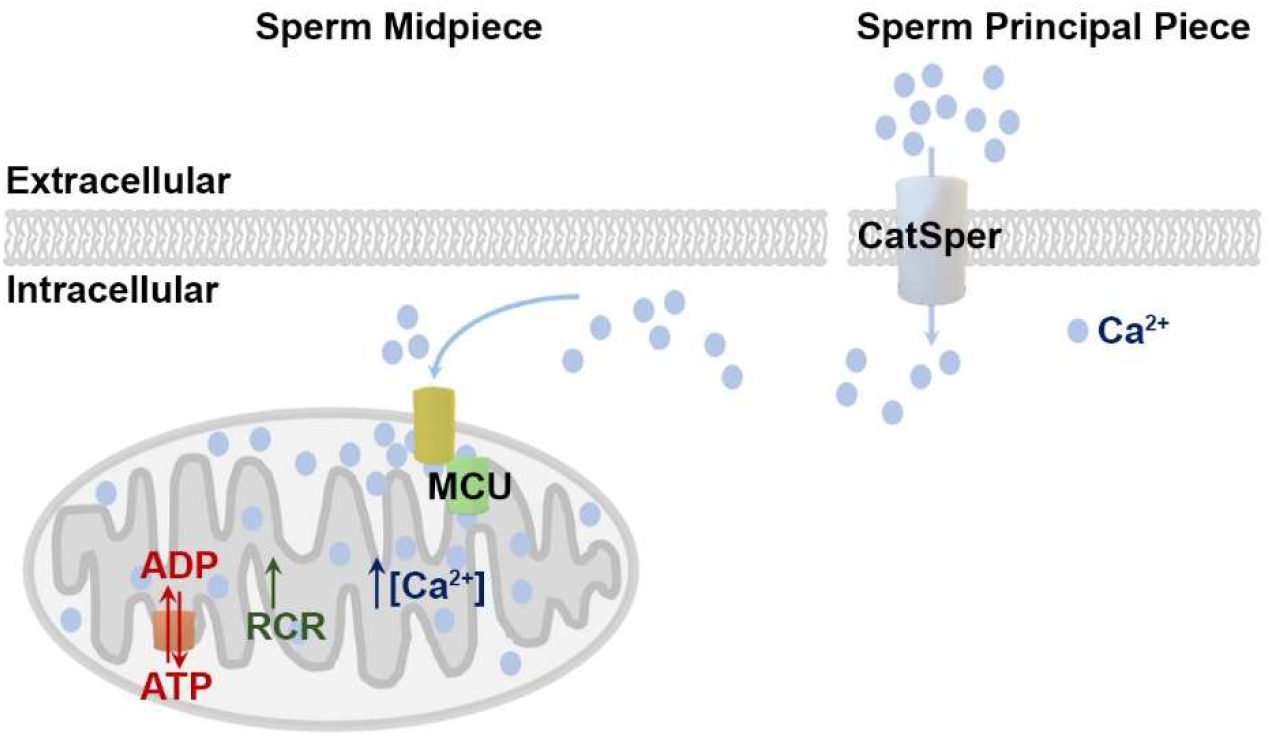
Proposed model for a new role of sperm mitochondria in capacitation/hyperactivation processes. CatSper activation during capacitation determines Ca^2+^ influx and the increase in cytoplasmic [Ca^2+^], that starts in the sperm principal piece, propagates through the midpiece, and reaches the head in a few seconds (20). Our data shows that the Ca^2+^ propagation through the midpiece leads to an increase in mitochondrial Ca^2+^ mediated by MCU. The increase in mitochondrial Ca^2+^ translates into an increase in mitochondrial efficiency that in turn promotes hyperactivation and *in-vitro* fertilization. Other molecular mechanisms involved in this pathway (e. g. possible role of the redundant nuclear envelop) need to be elucidated.

The role of mitochondria in mammalian sperm function, quality, and fertilization ability has been intensely debated for several years (15, 17, 18, 29–31). The contribution of mitochondria to sperm bioenergetics is unclear, and the source of the ATP used for sperm motility and hyperactivation has been long debated. The studies carried out in several species have provided different or conflicting results. In mouse sperm, glycolysis-produced ATP is sufficient to sustain progressive motility (15, 32). Conversely, in human sperm, a strong correlation has been noted between mitochondrial functionality and sperm motility or overall quality (33–35). Therefore, one idea is that sperm have versatile metabolism, allowing them to use species or environment dependent mechanisms for energy production (33, 36).

Here, we measured mitochondrial activity in NC and CAP mouse sperm by two independent methods. First, we used HRR to measure mitochondrial oxygen consumption in intact and motile sperm cells (21, 37, 38). During capacitation, mitochondrial coupling efficiency increased, indicating that mitochondrial electron transport chain is more closely coupled to ADP phosphorylation in CAP sperm than in NC sperm. Additionally, we noted an increase in RCR in CAP sperm, suggesting that mitochondrial function improved to increase ATP production, or to maintain the elevated MMP. However, the reserve respiratory capacity, which reflects the capacity of sperm to respond to energy demands, was not modified in CAP sperm, suggesting that sperm operate close to their bioenergetic limit (39). As a second method to study mitochondrial function, we used the voltage-sensitive dye TMRM (26) to measure sperm MMP both at the sperm population, and single cell level (23, 24), observing an increase in MMP and mitochondria function (40). Therefore, we conclude that sperm mitochondrial function increases during capacitation.

Our studies have a significant advantage over previous studies that also support a role of mitochondria in sperm capacitation, that is the fact that we conducted our experiments in intact and motile sperm, while earlier studies have used non-physiological conditions (permeabilized and/or non-motile sperm) (16–19, 31, 41).

The experiments with the inhibitors of mitochondrial function indicated that mitochondrial activity is not necessary for two features of capacitation – AR and tyrosine phosphorylation– but is contributing significantly to sperm hyperactivation and fertilization ability. These findings are consistent with observations that sperm from CatSper KO mice fail to hyperactivate and fertilize, but the AR and tyrosine phosphorylation are unaffected (42). Additionally, these findings are consistent with our observation that sperm from CatSper KO mice did not have an increase in RCR or MMP in capacitating conditions. Although the downstream targets of the CatSper-mediated cytoplasmic [Ca^2+^] increase have not been fully characterized (9, 10), sperm from CatSper KO have decreased NADH levels in the midpiece and a deficit in ATP production (8, 43). It is possible that the tail-to-head propagation of Ca^2+^ initiated by CatSper activation triggers an increase in [NAD] and may regulate ATP homeostasis (20). Moreover, our finding that CatSper channels are required for increased mitochondrial [Ca^2+^] in CAP sperm adds a new role for the sperm-specific Ca^2+^ channel.

Several lines of evidence suggest that, at least in somatic cells, regulated Ca^2+^ entry into mitochondria increases the efficiency of oxidative respiration (44, 45). Mitochondrial [Ca^2+^] is considered a central regulator of oxidative phosphorylation by mediating NADH production and controlling activity of pyruvate dehydrogenase, isocitrate dehydrogenase, and α-ketoglutarate dehydrogenase (11, 46). Mitochondrial [Ca^2+^] also plays an important role in regulating ATP synthase (46, 47) and can trigger the release of pro-apoptotic agents by the mitochondria (48). However, the precise role of Ca^2+^ in sperm mitochondria is still under debate (14). Some evidence suggests that maintenance of mitochondrial Ca^2+^ homoeostasis is essential for motility regulation in human sperm (49) and bovine sperm (50, 51). Whereas in contrast, other studies reported that sperm mitochondrial [Ca^2+^] is unaltered by mitochondrial uncoupling (52) and in bulls, mitochondrial activity in hyperactivated sperm appears not to be regulated by [Ca^2+^] (53). Thus, the role of Ca^2+^ in sperm may be species-specific.

Our data are consistent with findings that, in somatic cells, mitochondrial Ca^2+^ uptake in mouse sperm occurs through the MCU, which is known to control intracellular Ca^2+^ signals, cell metabolism, and cell survival (12, 54). Specifically, we found that the MCU inhibitor Ru360 (55) decreased mitochondrial [Ca^2+^] in mouse sperm during capacitation and inhibited sperm hyperactivation. These results agree with previous results showing that mitochondrial Ca^2+^ contributes to motility regulation in human sperm (49) and capacitation in bovine sperm (49, 51). Proteomic studies have confirmed that human sperm possess both MCU and MCU regulator 1 (14, 56), so this protein may be required for mitochondrial Ca^2+^ uptake in human sperm as well. MCU has a low affinity for Ca^2+^ uptake (12, 27), leading some to speculate that Ca^2+^ transfer into mitochondria occurs at highly specialized regions of close contact between mitochondria and endoplasmic reticulum called mitochondria-associated membranes (27, 57–59). Some of the molecular components of mitochondria-associated membranes are present in sperm, distributed in the acrosome and at the sperm neck an anterior midpiece. Endoplasmic reticulum in this location is referred as the redundant nuclear envelope (53, 60, 61). Further studies are needed to determine whether these sites of contact between the redundant nuclear envelope and mitochondria participate in the increase of mitochondrial [Ca^2+^] during capacitation.

In conclusion, our results show that mitochondrial activity (i.e., oxygen consumption, generation of the electrochemical gradient, ATP/ADP exchange) increases during sperm capacitation and this increase in activity contributes to sperm hyperactivation and sperm fertilization. The increase in mitochondrial coupling efficiency is consistent with Ca^2+^ influx through CatSper channels and Ca^2+^ entry into the mitochondria through the MCU. This increase in mitochondrial Ca^2+^ could translate either into an increase in ATP production due to the Ca^2+^ dependency of many of the mitochondrial enzymes or play a role in shaping intracellular calcium signals. Both mechanisms, acting independently or together, would be relevant.to achieve sperm hyperactivation. Our discovery of new mechanisms that explain sperm function may lead to the use of new molecules for fertility treatments and male contraception.

## Materials and methods

### Animals and ethics statement

All experimental procedures were approved by the Comisión honoraria de experimentación animal (CHEA), Uruguay, and by the Animals studies committee of Washington University (St. Louis, MO); and performed following the National Institute of Health Guide for the Care and Use of Laboratory Animals.

Animals were kept under a 12/12 hours dark/light cycle at a constant temperature of 22 ± 2°C with free access to food and water. Wild-type sperm cells were obtained from male mice from two different strains, CB6F1/J mice (80-120 days old) in Montevideo-Uruguay and C57BL/6 male (60-90 days old) in Saint Louis, MO, USA. CatSper1 knock-out mice (CatSper KO) were obtained from Jackson Laboratory. Oocytes were obtained from CB6F1/J female mice (4-8 weeks old). Mice were sacrificed via cervical dislocation.

### Sperm collection and motility analysis

Motility analysis was performed in parallel in two laboratories. In Montevideo-Uruguay (UdelaR), sperm was collected from cauda epididymis in TYH media (NaCl 119.3 mM, KCl 4.7 mM, CaCl_2_.2H_2_O 1.71 mM, KH_2_PO_4_ 1.2 mM, MgSO_4_.7H_2_O 1.2 mM, NaHCO_3_ 25.1 mM, Glucose 5.56 mM, Sodium pyruvate 0.51 mM, Phenol red (1%) 0.0006% supplemented with 4 mg/mL Bovine serum albumin) and incubated for 60 minutes at 37 °C and 5% CO2 to induce capacitation. Sperm suspensions were loaded into pre-warmed sperm counting chambers (depth 20 μm) (DRM-600, Millennium Sciences, Inc. CELL-VU^®^, NY) and placed on a microscope stage at 37 °C. Sperm motility was examined using CASA system (SCA6 Evolution, Microptic, Barcelona, Spain). The microscope used was Nikon (Japan) Eclipse E200 with phase contrast 100X equipped with Basler (Germany) acA780-75gc camera. The default settings included the following: frames acquired: 30; frame rate: 60 Hz; head size: 5-70 μm^2^. Sperm with hyperactivated motility were sorted using the following parameters: VCL>182 μm/s y LIN<50%, STR>57%. At least 500 sperm were analyzed in each experiment. In Saint Louis, MO, USA (Washington University) sperm was collected from cauda epididymis, incubated in NC HS at pH 7.4 for 15-20 minutes at 37 °C. After this time, the motile fraction of the sample was removed from the tube and split based on conditions to be tested. To achieve capacitation, sperm were incubated for 90 minutes at 37 °C in HS medium with 5 mg/ml Bovine serum albumin (BSA), and 15 mM NaHCO3 added (CAP HS). NC sperm were incubated for 90 °C at 37°C in HS without BSA and NaHCO3 (Chavez et al 2014). HS NC media (in mM): 135 NaCl, 5 KCl, 2 CaCl_2_, 1 MgSO_4_, 20 HEPES, 5 glucose, 10 lactic acid, 1 Na-pyruvate in pH 7.4 or as indicated. Drugs or inhibitors were added in NC or CAP HS media. For motility tests, 3 μl of the sample were placed into a 20 μm Leja standard count 4 chamber slide, pre-warmed at 37°C, and a minimum of 200 cells were counted. CASA analysis was performed with a Hamilton–Thorne digital image analyzer (HTR-CEROS II v.1.7; Hamilton–Thorne Research, Beverly, MA, United States). CASA settings used for the analysis were: objective Zeiss 10XNH; 30 frames were acquired at 60 Hz; camera exposure: 8 ms; camera gain: 300; integrated time: 500 ms; elongation max%: 100; elongation min%: 1; head brightness min 170; head size max: 50 μm^2^; head size min: 5 μm^2^; static tail filter: false; tail brightness min: 70; tail brightness auto offset: 8; tail brightness mode: manual; progressive STR (%): 80; progressive VAP (μm/s): 25. The criteria used to define hyperactivated sperm was: curvilinear velocity (VCL) >150 μm/s, lateral head displacement (ALH) >7.0 μm, and linearity coefficient (LIN).

### Acrosome reaction (AR) evaluation

AR status was evaluated both in NC and CAP sperm samples. Induced AR was obtained by incubating the cells with 10 μM of the calcium ionophore A23187 for 30 minutes prior the end of the capacitation process. Mitochondrial inhibitors were added to CAP media with and without A23187, and 2.5 μM Antimycin A (AA) (Sigma Aldrich, St. Louis, MO) or 2.5 μM Carbonyl cyanide-4-(trifluoromethoxy) phenylhydrazone (FCCP; Sigma-Aldrich). For all conditions 15 μl samples were placed onto glass slides, fixed in 4% paraformaldehyde for 30 minutes and washed twice with phosphate buffered saline (PBS).

After washing, slides were incubated with 0.22% Coomassie stain (Coomassie Blue G-250; Thermo Scientific, Massachusetts), 50% methanol, 10% glacial acetic acid, 40% water for 2 minutes. Excess dye was removed by washing thoroughly using distilled water. Slides were air-dried and coverslips were placed on slides using mounting medium at room temperature. Stained sperm were examined under bright field microscopy at 400X (Nikon E100, Japan) to verify the percentage of sperm that had undergone AR. A minimum of 200 sperm was evaluated in each experiment.

### In Vitro Fertilization (IVF) protocol

CB6F1/J female mice (4-8 weeks old) were super-ovulated using intraperitoneal administration of 5UI Pregnant Mare Serum Gonadotropin (PMSG) (Syntex, Argentina) followed by 5UI human Chorionic Gonadotropin (hCG) (Intervet, Netherlands) 48 hours later. After 12-15 hours of hCG injection, female mice were sacrificed, and the oocyte-cumulus complex was isolated in 250 mL of TYH. Fertilization wells containing 30-50 eggs were inseminated with sperm (final concentration of 2.5 × 106 cells/ml) that had been CAP for 1 hour with or without mitochondria inhibitors (AA or FCCP) depending on the experimental conditions, washed twice by centrifugation, and resuspended in TYH. After 3 hours of insemination, eggs were washed and left in fresh media. To assess fertilization, eggs were evaluated after 24 hours postinsemination, and two-cell stage embryos were counted.

### High-Resolution Respirometry and Respiration Control Ratio

Sperm cells oxygen consumption was determined by HRR. HRR integrates highly sensitive oxygraphs (Oxygraph-2 K; Oroboros Instruments GmbH, Innsbruck, Austria) with software (DatLab, version 4.2; Oroboros Instruments GmbH) that presents respiration in terms of oxygen rate (pmol O2/106 cells/sec). Basal oxygen consumption was measured for 10 minutes, then 2 μg/ml Oligomycin (Sigma-Aldrich, St. Louis, MO) were added to the chamber to block mitochondrial ATP synthase. Maximal respiration was obtained by subsequent 0.5 μM stepwise of FCCP. Finally, 2.5 μM Antimycin A (Sigma-Aldrich) was added to distinguish mitochondrial from residual (non-mitochondrial respiration) oxygen consumption. A total of 15 million sperms cells/ml per condition was evaluated. Stirring speed was set to 750 rpm. For each sperm sample **(Figure 1A)**, we measured mitochondrial basal respiration rate [1]. By subtracting the oligomycin-resistant respiration rate [2] from the basal respiration rate, we calculated the oxygen consumption rate linked to ATP synthesis [1-2]. We also measured maximal respiration rate in the presence of FCCP [3]. Finally, we measured the non-mitochondrial respiration rate [4], which we subtracted from all the other values. From these measurements, we calculated 1) Coupling efficiency [2/1] (ratio between respiration linked to ATP synthesis and basal respiration), RCR (ratio between maximum and oligomycin-resistant respiration rates) [3/[1-2]], and spare respiratory capacity (ratio between the maximum and basal respiration rates) [3/1]. It is important to note that the resulting values were internally normalized in each sample and were independent of cell number, protein mass, and viability (22, 39).

### Mitochondrial Membrane Potential measurements

Flow cytometry: Sperm collected from cauda epididymis was allowed to swim-out in TYH CAP media for 5-10 minutes. Tissue was extracted and the suspended cells were split into two samples with a final concentration of 106 cells/ml. One of the samples was treated immediately for 25 minutes at 37°C with 300 nmol/L TMRM (Sigma-Aldrich Inc., St Louis, MO, USA) and was considered as our (control) NC condition. The second sample was incubated for 90 minutes at 37°C and 5% CO_2_ to induce capacitation, and TMRM was added 25 minutes before finishing the incubation period. TMRM was washed from both conditions by centrifuging the samples at 400 g and resuspended in 400 ml of PBS. Half of the washed sample (500 μl) was analyzed by flow cytometry and the remaining (500 μl) was incubated with 20 μM FCCP for 15 minutes and analyzed. Values obtained with FCCP were used as Fmin to normalize the samples fluorescence. We used the Geo Mean fluorescence values of FCCP at each experiment as minimum fluorescence to normalized fluorescence of M1 and M2 and discard differences related to the loading of TMRM for each experiment.

Flow cytometry analysis was performed using a FACS Calibur flow cytometer (Becton Dickinson, San Jose, CA). Cellular size and granularity were analyzed in the forward and side scatter (FSC-H and SSC-H), respectively. For each sample, 30000 single events were recorded in the forward light scatter/side light scatter dot plot. A gate was used to separate sperm from debris. TMRM was detected using a 585/42 nm bandwidth filter (FL-2). Samples were analyzed with Cell Quest software. Two populations of cells (M1 and M2, **Figure 1-C**) with different MMP were consistently obtained. Based on, MMP measurements after capacitation for both populations by our laboratory, previously described methodology by Uribe P. et al (23), and similar results published using TMRM in sperm by Yang Q. et al (62), we analyzed the population with higher MMP (M2).

Single Cell MMP measurements by Confocal Imaging: Sperm cells were collected from cauda epididymis. Sperm were obtained by Swim-out in HS NC or CAP media for 10-15 minutes. NC samples were incubated in NC HS media, and CAP samples in CAP HS media, for 90 minutes at 37°C. At 60 minutes of incubation 200-300 nmol/L TMRM was added to all conditions and incubated for 30 minutes at 37°C. TMRM was washed from all samples by centrifugation and cells were resuspended in NC HS media and allowed to attach for 5-10 minutes to pre-coated coverslips with Poly-L-lysine (0.1%). Basal TMRM fluorescence for all the different conditions was measured, before adding 20 μM FCCP to assess Fmin used to normalize initial values.

Fluorescence values from different animals and different conditions were adjusted to the same laser intensity and gain voltage. Recordings were performed by using a confocal microscope (Leica SP8) and LAS X 3.5.2.18963 software equipped with an ACS APO 40x, 1.15 Numerical aperture objective, and oil-immersion objective. The parameters for the acquisition were: “xyt” images, 1024×1024 pixels, pixel size 0.269 μm, pinhole of 600.0 μm, 400 Hz unidirectional sampling, frames were obtained every 30 seconds. Excitation was performed by 543-nm laser and emitted fluorescence was measured at 556 nm – 600 nm. All experiments were performed at room temperature (22–23°C). Fluorescence was measured only in regions of interest (ROI) corresponding to the sperm midpiece, using ImageJ software (version 1.48). Data was analyzed using pclamp 10 and Sigmaplot 12.

### Fluo-5N and Mitochondrial markers co-localization Images

Sperm was collected by swim-up in HS media from wild-type mice (C57Bl6) or Acr-eGFP+Su9-Red2 transgenic mice, loaded with 2-4 μM Fluo-5N and incubated with 200 nM MitoTracker^^®^^ Red CMXRos – M7512 (Invitrogen, USA). To load Fluo-5N, cells were incubated at 37°C for 45-60 minutes in HS with 2 μM Fluo-5N AM and 0.05-0.1% Pluronic Acid F-127. MitoTracker was added 20-30 minutes before Fluo-5N loading was finished. After, sperm was centrifuged, resuspended in HS NC media, and allowed to attach for 5-10 minutes to pre-coated coverslips with Poly-L-lysine (0.1%). Images were acquired using a confocal microscope (Leica SP8) and LAS X 3.5.2.18963 software equipped with an ACS APO 63x, 1.15 Numerical aperture objective, oil-immersion objective. Images dimensions were 2048×2048 pixels. Excitation was provided by 488 nm and 543 nm lasers and emitted fluorescence was measured at 500–530 and >560 nm, for Fluo-5N and MitoTracker respectively. All experiments were performed at room temperature (22–23°C). ImageJ software (version 1.48) was used to analyze the images.

### Mitochondrial Ca^2+^ measurements

After swim-up in HS NC or CAP media, motile cells were incubated with 2-4 μM Fluo-5N AM and 0.05% Pluronic F-127 at 37°C for 60-90 minutes. After loading, sperm cells were centrifuge at 1500-2000 rpm for 5-10 minutes and resuspended in the corresponding media. Sperm was allowed to attach to Poly-L-lysine (0.1%) coated coverslips placed on the recording chamber floor for 5 min. A local perfusion device with an estimate exchange time of 10 seconds was used to applied various test solutions. Calcium signals were recorded with a Leica AF 6000LX system with a Leica DMi8000 inverted microscope and an Andor-Zyla-VCS04494 camera. A halogen lamp was used with a 488 +/-20 nm excitation filter and a 530 +/-20 nm emission filter. A 63X objective (HC PL FluoTar L 63X/0.70 Dry) air objective was used. Data was collected with Leica LasX 2.0.014332 software. Acquisition parameters were: 20 ms exposure time, 4×4 binning, 1024 x 1024 pixels resolution. Whole images were collected every 10 seconds. LAS X, ImageJ, Clampfit 10 (Molecular Devices), and SigmaPlot 12 were used to analyze data.

ROI were selected in the sperm midpiece. Reference Fluo-5N fluorescence was measured at the beginning of the experiments (F_Ref_), for all the conditions. To compared F_Ref_ levels across different animals and experimental conditions, we normalized Fluo-5N fluorescence using Fmin and Fmax obtained from each cell. F_min_ was obtained by perfusing cells with 0 mM Ca^2+^ + 2 mM EGTA and 2-5 μM Ionomycin. F_max_ was obtained by perfusing the cells with 2 mM Ca^2+^ + Ionomycin. Experiments were performed at room temperature.

### Statistical analysis

Sigmaplot version 12.0 (Systat Software Inc.), and GraphPad Prism version 6.01 for Windows (GraphPad Software, La Jolla California USA http://www.graphpad.com) were used for all statistical analysis. An unpaired Student’s t-test was used to compare independent samples, and a paired t-test was used to compare data in studies performed in the same sample. Data are expressed as the mean ± SD. P-value < 0.05 was considered statistically significant.

## Acknowledgements

We thank Dr. Deborah Frank for the critical review of the manuscript, as well as Dr. Emily Hunter and Nathan Pomper for the editing assistance provided through InPrint: A Scientific Editing Network at Washington University in St. Louis. We thank Dr. Lis Puga Molina for her assistance with the CASA experiments.

## Reference

1. Chang MC (1951) Fertilizing Capacity of Spermatozoa Deposited into the Fallopian Tubes. Nature 168(4277):697–698.

2. Austin CR (1952) The Capacitation of the Mammalian Sperm. Nature 170(4321):326–326.

3. Gervasi MG & Visconti PE (2016) Chang’s meaning of capacitation: A molecular perspective. Mol Reprod Dev 83(10):860–874.

4. Correia J, Michelangeli F, & Publicover S (2015) Regulation and roles of Ca2+ stores in human sperm. Reproduction 150(2):R65–76.

5. Ruknudin A & Silver IA (1990) Ca2+ uptake during capacitation of mouse spermatozoa and the effect of an anion transport inhibitor on Ca2+ uptake. Mol Reprod Dev 26(1):63–68.

6. Lishko PV & Mannowetz N (2018) CatSper: A Unique Calcium Channel of the Sperm Flagellum. Current opinion in physiology 2:109–113.

7. Kirichok Y, Navarro B, & Clapham DE (2006) Whole-cell patch-clamp measurements of spermatozoa reveal an alkaline-activated Ca2+ channel. Nature 439(7077):737–740.

8. Ren D, et al. (2001) A sperm ion channel required for sperm motility and male fertility. Nature 413(6856):603–609.

9. Vyklicka L & Lishko PV (2020) Dissecting the signaling pathways involved in the function of sperm flagellum. Curr Opin Cell Biol 63:154–161.

10. Rahban R & Nef S (2020) CatSper: The complex main gate of calcium entry in mammalian spermatozoa. Molecular and cellular endocrinology 518:110951.

11. McCormack JG, Halestrap AP, & Denton RM (1990) Role of calcium ions in regulation of mammalian intramitochondrial metabolism. Physiological reviews 70(2):391–425.

12. Rizzuto R, De Stefani D, Raffaello A, & Mammucari C (2012) Mitochondria as sensors and regulators of calcium signalling. Nature reviews. Molecular cell biology 13(9):566–578.

13. Bravo-Sagua R, et al. (2017) Calcium Transport and Signaling in Mitochondria. Comprehensive Physiology 7(2):623–634.

14. Amaral A, Lourenco B, Marques M, & Ramalho-Santos J (2013) Mitochondria functionality and sperm quality. Reproduction 146(5):R163–174.

15. Miki K, et al. (2004) Glyceraldehyde 3-phosphate dehydrogenase-S, a sperm-specific glycolytic enzyme, is required for sperm motility and male fertility. Proceedings of the National Academy of Sciences of the United States of America 101(47):16501–16506.

16. Boell EJ (1985) Oxygen consumption of mouse sperm and its relationship to capacitation. The Journal of experimental zoology 234(1):105–116.

17. Stendardi A, et al. (2011) Evaluation of mitochondrial respiratory efficiency during in vitro capacitation of human spermatozoa. International journal of andrology 34(3):247–255.

18. Ferramosca A & Zara V (2014) Bioenergetics of mammalian sperm capacitation. BioMed research international 2014:902953.

19. Balbach M, et al. (2020) Metabolic changes in mouse sperm during capacitationdagger. Biology of reproduction 103(4):791–801.

20. Xia J, Reigada D, Mitchell CH, & Ren D (2007) CATSPER channel-mediated Ca2+ entry into mouse sperm triggers a tail-to-head propagation. Biology of reproduction 77(3):551–559.

21. Cassina A, et al. (2015) Defective Human Sperm Cells Are Associated with Mitochondrial Dysfunction and Oxidant Production. Biology of reproduction 93(5):119.

22. Buckman JF & Reynolds IJ (2001) Spontaneous changes in mitochondrial membrane potential in cultured neurons. J Neurosci 21(14):5054–5065.

23. Uribe P, et al. (2017) Use of the fluorescent dye tetramethylrhodamine methyl ester perchlorate for mitochondrial membrane potential assessment in human spermatozoa. Andrologia 49(9).

24. Creed S & McKenzie M (2019) Measurement of Mitochondrial Membrane Potential with the Fluorescent Dye Tetramethylrhodamine Methyl Ester (TMRM). Methods Mol Biol 1928:69–76.

25. Kabbara AA & Allen DG (2001) The use of the indicator fluo-5N to measure sarcoplasmic reticulum calcium in single muscle fibres of the cane toad. The Journal of physiology 534(Pt 1):87–97.

26. Visconti, et al. (1995) Capacitation of mouse spermatozoa. I. Correlation between the capacitation state and protein tyrosine phosphorylation. Development.

27. Pathak T & Trebak M (2018) Mitochondrial Ca(2+) signaling. Pharmacology & therapeutics 192:112–123.

28. Kirichok Y, Krapivinsky G, & Clapham DE (2004) The mitochondrial calcium uniporter is a highly selective ion channel. Nature 427(6972):360–364.

29. Nascimento JM, et al. (2008) Comparison of glycolysis and oxidative phosphorylation as energy sources for mammalian sperm motility, using the combination of fluorescence imaging, laser tweezers, and real-time automated tracking and trapping. J Cell Physiol 217(3):745–751.

30. Ramalho-Santos J, et al. (2009) Mitochondrial functionality in reproduction: from gonads and gametes to embryos and embryonic stem cells. Human reproduction update 15(5):553–572.

31. Ferramosca A, Focarelli R, Piomboni P, Coppola L, & Zara V (2008) Oxygen uptake by mitochondria in demembranated human spermatozoa: a reliable tool for the evaluation of sperm respiratory efficiency. International journal of andrology 31(3):337–345.

32. Mukai C & Okuno M (2004) Glycolysis plays a major role for adenosine triphosphate supplementation in mouse sperm flagellar movement. Biology of reproduction 71(2):540–547.

33. Ruiz-Pesini E, Diez-Sanchez C, Lopez-Perez MJ, & Enriquez JA (2007) The role of the mitochondrion in sperm function: Is there a place for oxidative phosphorylation or is this a purely glycolytic process? Curr Top Dev Biol 77:3–19.

34. Ruiz-Pesini E, et al. (1998) Correlation of sperm motility with mitochondrial enzymatic activities. Clin Chem 44(8):1616–1620.

35. Gallon F, Marchetti C, Jouy N, & Marchetti P (2006) The functionality of mitochondria differentiates human spermatozoa with high and low fertilizing capability. Fertility and sterility 86(5):1526–1530.

36. Storey BT (2008) Mammalian sperm metabolism: oxygen and sugar, friend and foe. The International journal of developmental biology 52(5-6):427–437.

37. Darr CR, Cortopassi GA, Datta S, Varner DD, & Meyers SA (2016) Mitochondrial oxygen consumption is a unique indicator of stallion spermatozoal health and varies with cryopreservation media. Theriogenology 86(5):1382–1392.

38. Moraes CR & Meyers S (2018) The sperm mitochondrion: Organelle of many functions. Anim Reprod Sci 194:71–80.

39. Brand MD & Nicholls DG (2011) Assessing mitochondrial dysfunction in cells. The Biochemical journal 435(2):297–312.

40. Moscatelli N, et al. (2017) Single-cell-based evaluation of sperm progressive motility via fluorescent assessment of mitochondria membrane potential. Sci Rep-Uk 7.

41. Balbach M, Buck J, & Levin LR (2020) Using an Extracellular Flux Analyzer to Measure Changes in Glycolysis and Oxidative Phosphorylation during Mouse Sperm Capacitation. Journal of visualized experiments: JoVE (155).

42. Qi H, et al. (2007) All four CatSper ion channel proteins are required for male fertility and sperm cell hyperactivated motility. Proceedings of the National Academy of Sciences of the United States of America 104(4):1219–1223.

43. Quill TA, et al. (2003) Hyperactivated sperm motility driven by CatSper2 is required for fertilization. Proceedings of the National Academy of Sciences of the United States of America 100(25):14869–14874.

44. Evtodienko YV (2000) Sustained oscillations of transmembrane Ca2+ fluxes in mitochondria and their possible biological significance. Membrane Cell Biology.

45. Cortassa S, Aon MA, Marban E, Winslow RL, & O’Rourke B (2003) An integrated model of cardiac mitochondrial energy metabolism and calcium dynamics. Biophys J 84(4):2734–2755.

46. Mccormack JG & Denton RM (1993) The Role of Intramitochondrial Ca2+ in the Regulation of Oxidative-Phosphorylation in Mammalian-Tissues. Biochemical Society Transactions 21(3):793–799.

47. Territo PR, French SA, Dunleavy MC, Evans FJ, & Balaban RS (2001) Calcium activation of heart mitochondrial oxidative phosphorylation: rapid kinetics of mVO2, NADH, AND light scattering. The Journal of biological chemistry 276(4):2586–2599.

48. Szalai G, Krishnamurthy R, & Hajnoczky G (1999) Apoptosis driven by IP(3)-linked mitochondrial calcium signals. The EMBO journal 18(22):6349–6361.

49. Bravo A, et al. (2015) Effect of mitochondrial calcium uniporter blocking on human spermatozoa. Andrologia 47(6):662–668.

50. Meyers S, Bulkeley E, & Foutouhi A (2019) Sperm mitochondrial regulation in motility and fertility in horses. Reproduction in domestic animals = Zuchthygiene 54 Suppl 3:22–28.

51. Rodriguez PC, Satorre MM, & Beconi MT (2012) Effect of two intracellular calcium modulators on sperm motility and heparin-induced capacitation in cryopreserved bovine spermatozoa. Anim Reprod Sci 131(3-4):135–142.

52. Machado-Oliveira G, et al. (2008) Mobilisation of Ca2+ stores and flagellar regulation in human sperm by S-nitrosylation: a role for NO synthesised in the female reproductive tract. Development 135(22):3677–3686.

53. Ho HC & Suarez SS (2003) Characterization of the intracellular calcium store at the base of the sperm flagellum that regulates hyperactivated motility. Biology of reproduction 68(5):1590–1596.

54. Patron M, et al. (2013) The mitochondrial calcium uniporter (MCU): molecular identity and physiological roles. The Journal of biological chemistry 288(15):10750–10758.

55. Matlib MA, et al. (1998) Oxygen-bridged dinuclear ruthenium amine complex specifically inhibits Ca2+ uptake into mitochondria in vitro and in situ in single cardiac myocytes. The Journal of biological chemistry 273(17):10223–10231.

56. Wang H, et al. (2020) TRPC channels: Structure, function, regulation and recent advances in small molecular probes. Pharmacology & therapeutics 209:107497.

57. G. Csordás, A. P. Thomas, & Hajnóczky G (2001) Calcium Signal Transmission between Ryanodine Receptors and Mitochondria in Cardiac Muscle. Trends in Cardiovascular Medicine 11(7):269–275.

58. Moreau B, Straube S, Fisher RJ, Putney JW, Jr., & Parekh AB (2005) Ca2+-calmodulin-dependent facilitation and Ca2+ inactivation of Ca2+ release-activated Ca2+ channels. The Journal of biological chemistry 280(10):8776–8783.

59. Piomboni P, Focarelli R, Stendardi A, Ferramosca A, & Zara V (2012) The role of mitochondria in energy production for human sperm motility. International journal of andrology 35(2):109–124.

60. Ho K, Wolff CA, & Suarez SS (2009) CatSper-null mutant spermatozoa are unable to ascend beyond the oviductal reservoir. Reprod Fert Develop 21(2):345–350.

61. Dragileva E, Rubinstein S, & Breitbart H (1999) Intracellular Ca(2+)-Mg(2+)-ATPase regulates calcium influx and acrosomal exocytosis in bull and ram spermatozoa. Biology of reproduction 61(5):1226–1234.

62. Yang Q, et al. (2020) Ca(2+) ionophore A23187 inhibits ATP generation reducing mouse sperm motility and PKA-dependent phosphorylation. Tissue & cell 66:101381.

