## Supplementary Figures for "Increased mitochondrial activity upon CatSper channel activation is required for sperm capacitation"

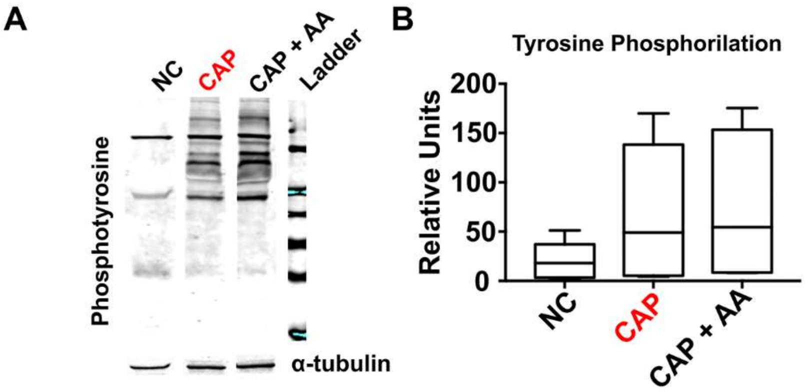


**Supplementary Figure 1. Tyrosine phosphorylation increases during capacitation and is not impaired by the mitochondrial inhibitor Antimycin A. A.** Representative western blots for tyrosine phosphorylated proteins. **B.** Protein levels in independent western blots were quantified by densitometry and normalized using tubulin as loading control. Quantification of bands is expressed as mean +/- SEM in units of density (n=6). Semen samples were centrifuged at 500 g and the pellet was resuspended in lysis buffer supplemented with protease and phosphatase inhibitors. Suspensions were sonicated for 5 seconds and then centrifuged at 14000 g for 10 minutes at 4 C. Protein concentration was determined by the Bradford method (63).

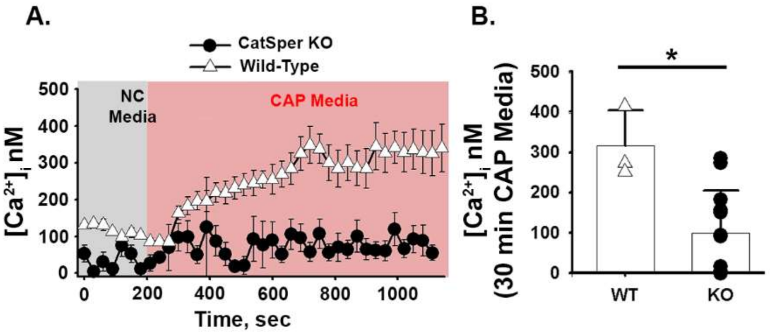


**Supplementary Figure 2. Intracellular calcium increases during Wt sperm capacitation but not in CatSper KO mice.** Ratiometric measurements with Fura 2 AM were used to calculate intracellular free calcium in WT and CatSper KO mice. Free swimming epididymal (Cauda) sperm cells were obtained using the swim-up technique in NC Media. Cells were loaded for 45 min with Fura-2 AM at a concentration of 4 μM. Before recordings, sperm was centrifuge and resuspended in the NC HS Media. **A**. Average traces of calculated intracellular calcium concentration $[Ca^{2+}]_i$ for WT and CatSper KO mice (REF calculation method). Recordings were started in HS Non-Capacitated and switched at 3-4 min to Capacitated HS Media. Bars are SEM. **B.** Graph. Average of $[Ca^{2+}]_i$ nM for WT and KO mice at 20-30 min of incubation in HS Capacitated Media. Values: for WT Average 315.32 nM +/- 88.67 SD, n=3. For CatSper KO Average: 99.01 nM +/- 104.90 SD, n=13.

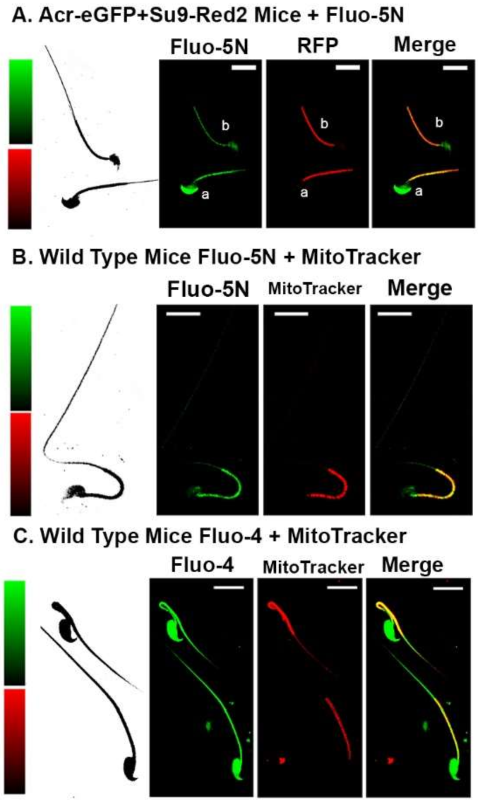


**Supplementary Figure 3. Fluo-5N fluorescence colocalizes with mitochondria markers in mouse sperm.** **A**. Acr-eGFP+Su9-Red2 transgenic mice expressing GFP in sperm acrosome and RFP in sperm mitochondria loaded with Fluo-5N AM. Acr-eGFP+Su9-Red2 loaded with Fluo-5N AM**.** Localization of RFP in the midpiece of the sperm. Merge of RFP and Fluo-5N showing co-localization in yellow. **B.** Wild-type C57BL6 mice sperm loaded with Fluo-5N AM and MitoTracker AM. Merged image of Fluo-5N and MitoTracker. Colocalization showed in yellow. Middle and right panel show sperm cells without and with acrosome respectively. **C.** Wild-type mice sperm loaded with Fluo-4 AM and MitoTracker AM. Merge of Fluo-4 and MitoTracker.

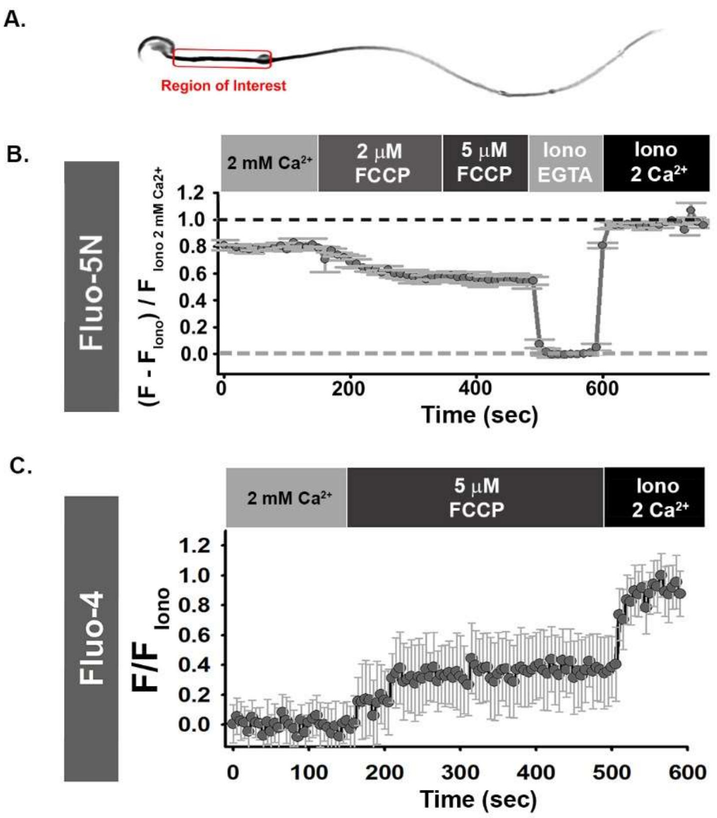


**Supplementary Figure 4. Effect of FCCP on sperm cells loaded with Fluo-5N AM or with Fluo-4 AM**. A Scheme indicating that fluorescence was measured only in the sperm midpiece. **B.** Representative recording of Fluo-5N fluorescence under conditions indicated in the figure. **C.** Representative recording of Fluo-4 fluorescence under conditions indicated in the figure. Error bars represent SD. Fluo-5N fluorescence ratios were calculated as = $F_{Initial} - (F_{Iono\ EGTA}) / F_{Iono\ 2Ca2+}$. For Fluo-4 AM loading protocol, see Supplementary Figure 5.

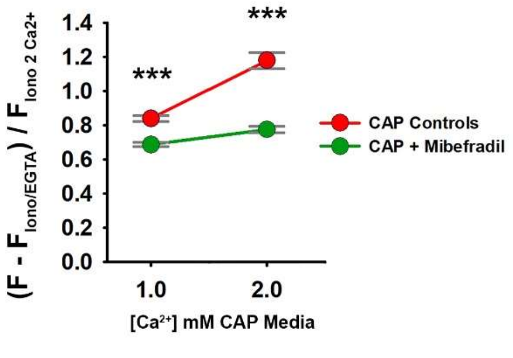


**Supplementary Figure 5. Mitochondrial [$Ca^{2+}$] increase during capacitation is reduced by the CatSper channel inhibitor Mibefradil.** Graph shows initial fluorescence values normalized to $F_{Iono\ EGTA}$ (0 mM $Ca^{2+}$ + Ionomycin + EGTA) / $F_{Iono2Ca2+}$ (Ionomycin + 2 mM $Ca^{2+}$). Recordings were obtained at 1 and 2 mM extracellular $Ca^{2+}$ for both conditions (CAP Control and CAP with 20 µM Mibefradil). Independent *t-test* were used to evaluate statistical significance between groups at the same extracellular [$Ca^{2+}$]. *** $P < 0.001$. Error Bars represent SD. n=42-138 from 3 mice for each condition.

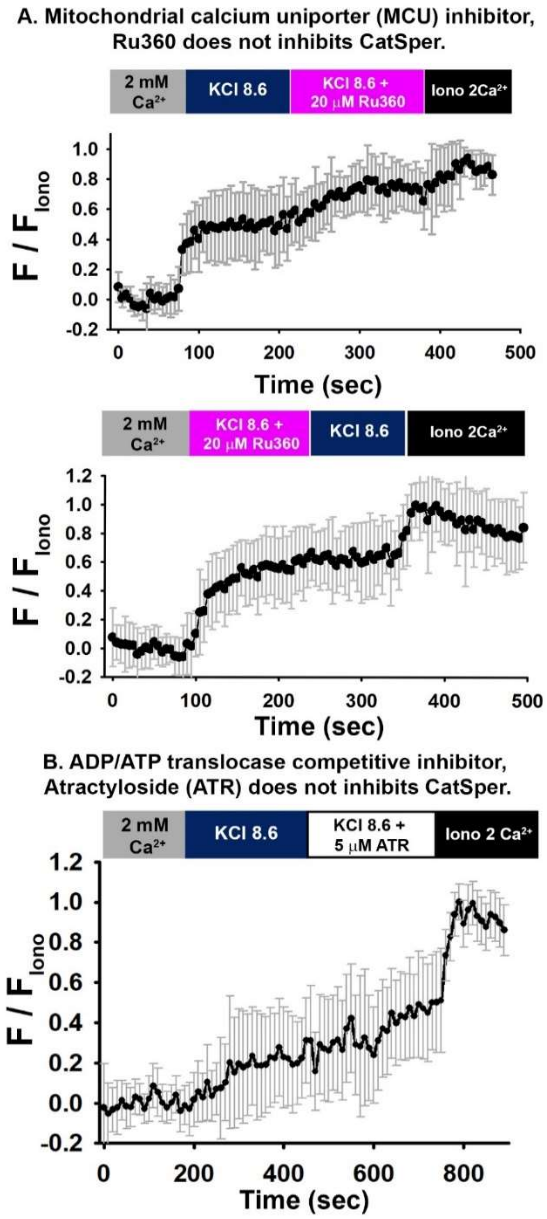


**Supplementary Figure 6. Ru360 and ATR have no effect on CatSper channels in CAP mouse sperm.** Sperm cells were loaded with Fluo-4 AM and CatSper activation was triggered by alkaline depolarization (50 mM KCl, pH 8.6). **A** Top: average traces + SD of Fluo-4 fluorescence recorded under conditions indicated in the fig. 20 μM Ru360 was added after stimulation of CatSper channels with alkaline depolarization. Bottom: 20 μM R360 was added simultaneously with 50 mM KCl pH 8.6. **B**. Average traces of Fluo-4 fluorescence recorded in the conditions indicated in the fig. 5 μM ATR was added after stimulation of CatSper channels with alkaline depolarization. Errors bars are SD. Protocol: After swim-up, motile cells were incubated with 2-4 μM Fluo-4 AM and 0.05% Pluronic F-127 in HS media at 37 °C for 90 minutes. Then, cells were centrifuge at 1500 rpm for 10 minutes and re-suspended in the HS media. Sperm were allowed to attach to laminin (1 mg/ml), Poly-L-lysine (0.1%) or cell-tak coated coverslips placed at the bottom of the recording chamber for 5 minutes. Ionomycin (5 μM) and HS 2 mM $Ca^{2+}$ were added at the end of the recordings as references of maximum fluorescence.
